# Characterization of Coxsackievirus A20 from a child with acute flaccid paralysis in Nigeria

**DOI:** 10.1101/399212

**Authors:** T.O.C. Faleye, M.O. Adewumi, O.T. Olayinka, J.A. Adeniji

**Affiliations:** Department of Virology, College of Medicine, Faculty of Basic Medical Sciences, University of Ibadan, Ibadan, Nigeria; Department of Microbiology, Faculty of Science, Ekiti State University, Ado-Ekiti, Nigeria; WHO National Polio Laboratory, University of Ibadan, Ibadan, Nigeria

## Abstract

In light of the ongoing cVDPV2 outbreak in Nigeria, we describe the draft genome of a CVA20 strain from a child with AFP. The non-structural region of this genome unambiguously unveiled the source of such regions in recombinant cVDPV2s (JX275140 and KX162716) found in Nigeria in 2008 and 2015, respectively.

## Introduction

Enteroviruses (EVs) belong to the genus Enterovirus, family Picornaviridae, order Picornavirales. The genus has 15 species and Poliovirus (the best characterized member of the genus) belongs to Species C. Several Coxsackie A viruses(e.g. CVA20) and some numbered EVs are nonpolio enterovirus member of Species C (NPEV-C).

In 2016, we (Adeniji et al., 2018) detected a CVA20 strain during our investigation of non-reproducible cytopathic effect (CPE) in L20B cell line. The virus produced CPE on initial inoculation into L20B cell line but the cytopathology was not reproducible on passage in L20B and RD cell lines (Adeniji et al., 2018). This phenomenon had been ascribed to nonspecific cytotoxicity, Adenoviruses and/or Reoviruses (WHO, 2011; Thorley and Roberts, 2016) but we provided evidence that enteroviruses could also contribute to the phenomenon (Adeniji et al., 2018). Further, considering the history of Nigeria and circulating Vaccine Derived Poliovirus 2s (cVDPV2s) (Burns et al., 2013),the ongoing cVDPV2 outbreak and the contributions of NPEV-Cs to emergence of cVDPV2, we attempted to establish the CVA20 virus in cell culture, sequence its genome and see what we can glean from it about recombinant cVDPV2s in Nigeria.

## The Study

Cell culture supernatant from the L20B tube that was inoculated with stool suspension from a child with AFP and initially scored positive for CPE in 2016 was recovered from storage at – 80°C. It was thenpassaged in HEp 2C cell line. By day seven, CPE developed and was reproducible on passage in HEp 2C cell line. After three freeze-thaw cycles, viral RNA was extracted from the isolate using Total RNA extraction kit (Jena Bioscience, Jena, Germany) following the manufacturer’s instructions. Complementary DNA (cDNA) was subsequently made using Random Hexamers with the SCRIPTcDNA synthesis kit (Jena Bioscience, Jena, Germany) as recommended by the manufacturer.

The genome of the isolate was amplified in overlapping fragments (2 - 3Kb each) using a blend of previously described (Nix et al., 2006,Oberste et al., 2006, Bessaud et al., 2008; 2016, Bailly et al., 2011,Arita et al., 2015) primers and Redload PCR kit (Jena Bioscience, Jena, Germany). Amplicons generated were pooled and shipped to a commercial facility (MR, Texas, USA) where library preparation and NextGen sequencing was done. Sequencing was done paired end for 300 cycles using the HiSeq system (Illumina). Assembly was done using the Kiki Assembler v0.0.9.

Two contigs were recovered spanning nucleotides 289 to 2,131 (MH785183; 5’-UTR to VP3) and 2,602 to 7,381 (MH785184; VP1 to 3’-UTR) relative to reference sequence NC_002058. A region of about 500nt (spanning the VP3/VP1 junction) that should join the contigs was missing despite being part of an amplicon (5’-UTR to VP1) generated by primers C004 (Bessaud et al., 2016) and AN88 (Nix et al., 2006) and present in the pool sequenced. In all, 88.9% of the genome of the CVA20 described in this study was recovered and the isolate confirmed as CVA20 by the enterovirus genotyping tool (Figure 1a). Phylogenetic analysis of the VP1 gene confirmed the isolate as CVA20 (Figure 2) and as the same virus described in (Adeniji et al., 2018). A BLAST search further confirmed the isolate as CVA20 (data not shown). It also showed that the P2 and P3 non-structural region of the virus was most similar to JX275140 and KX162716, respectively (data not shown). JX275140 and KX162716 are both recombinant cVDPV2s with non-structural region of unknown origin and were recovered in Nigeria. JX275140 was isolated in 2008 (Burns et al., 2013) while KX162716 was isolated in 2015 (Montmayeur et al., 2017).

**Figure 1:**
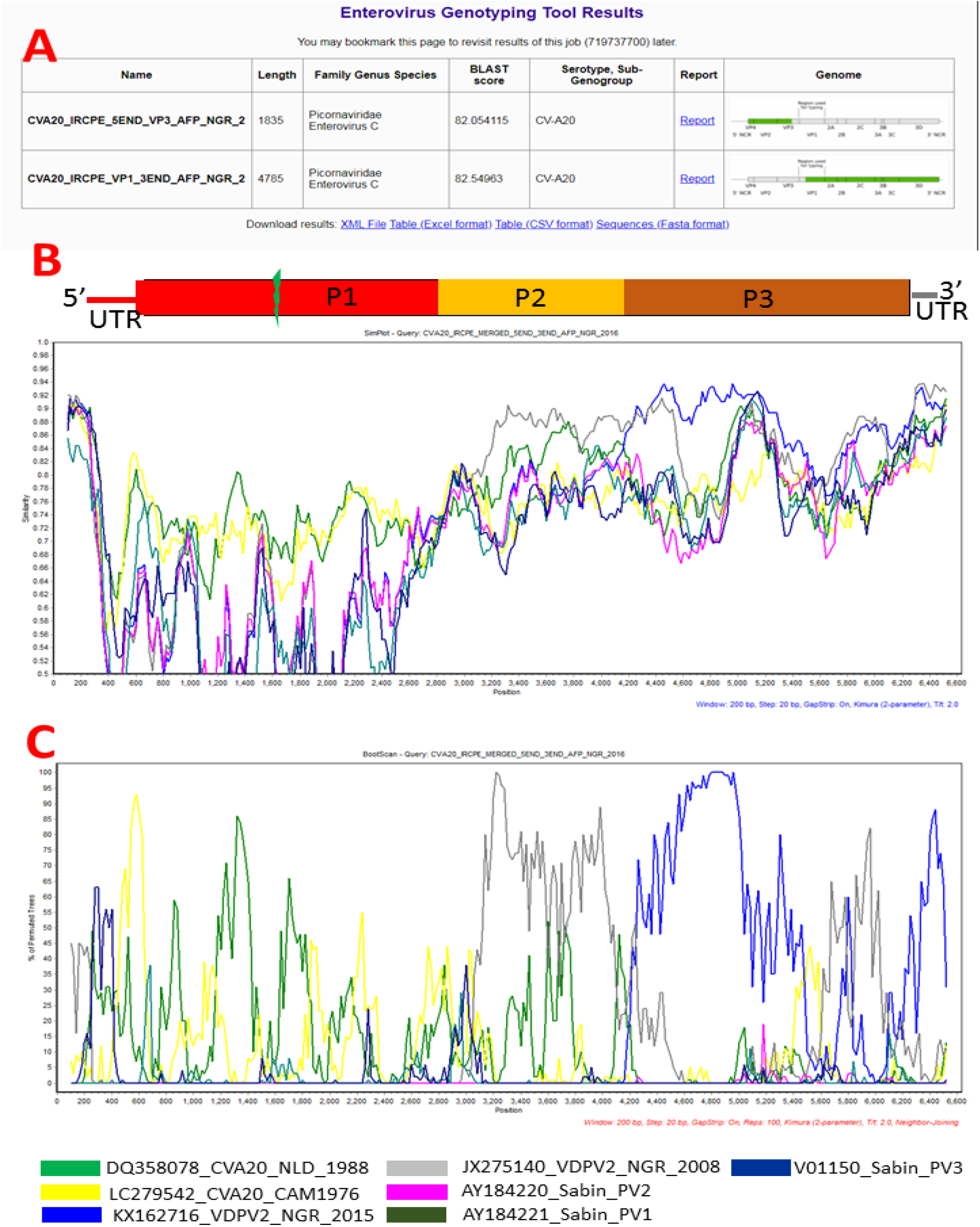
Enterovirus Genotyping Tool display of the genomic regions covered in the two different contigs (1a). Schematic representation of enterovirus genomic organisation and result of SimPlot (1b). Result of BootScan analysis (1c). Note: The green staggered line in the P1 region of the genome’s schematic representation (1b),denotes the fact that a 500nt window missing from the CVA20 genome was deleted from the alignment and consequently on both SimPlot and BootScan analysis.

**Figure 2:**
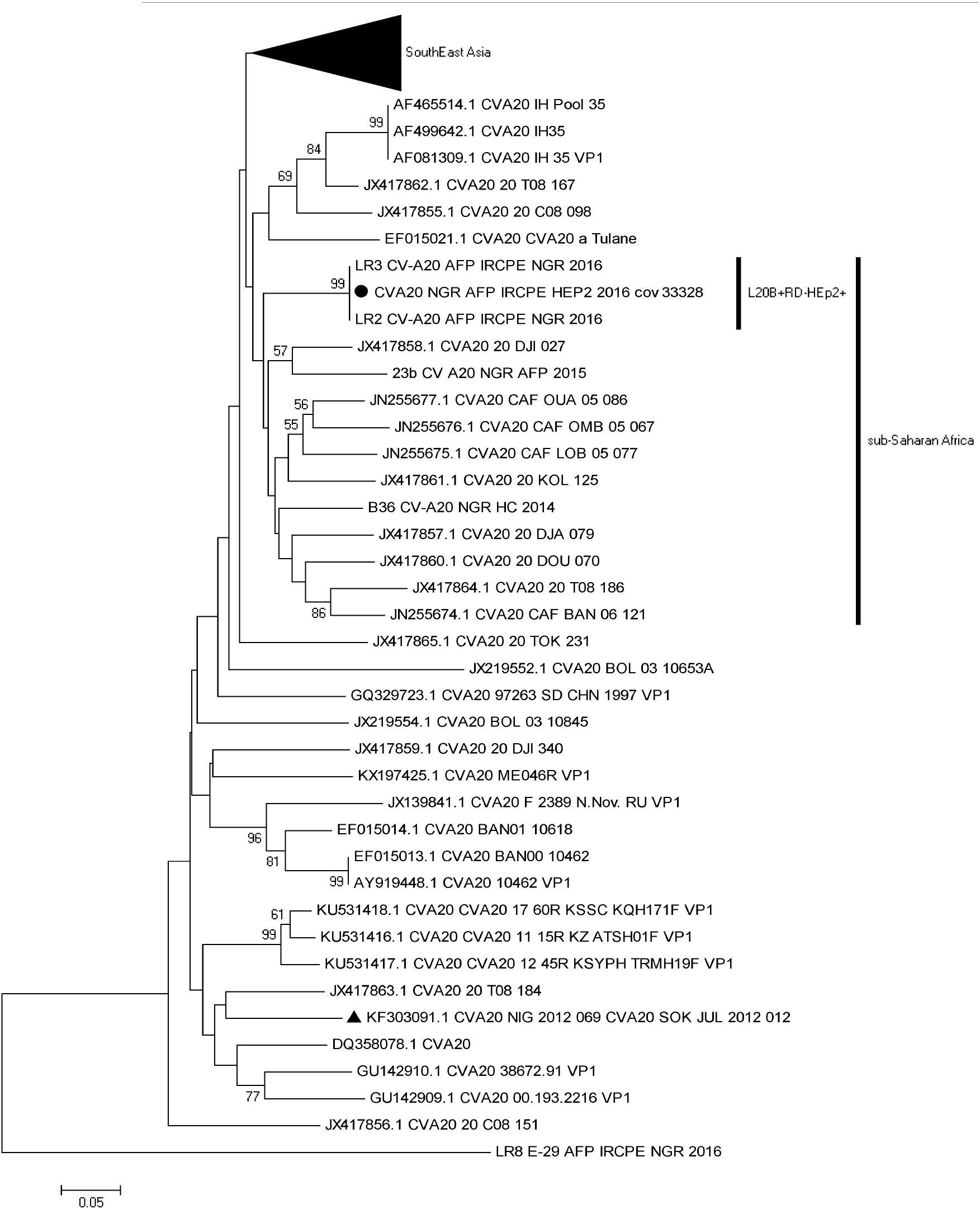
Phylogram of genetic relationship between VP1 nucleotide sequences of CVA20. The phylogenetic tree is based on an alignment of partial VP1 sequences. The newly sequenced strain is indicated with black circle. The strain recovered in Nigeria in 2012 from Sewage contaminated water is indicated with a black triangle. Bootstrap values are indicated if >50%.

To confirm the similarity between JX275140, KX162716 and the CVA20 described in this study, the sequences were aligned alongside reference CVA20s and Sabin PV1-3 strains. Alignment was done using the ClustalW programme in MEGA 5 software. Analysis for similarity and recombination was done using SimPlot version 3.5. Both SimPlot (Figure 1b) and Bootscan (Figure 1c) analysis confirmed the isolate as CVA20 in the P1 region. They also confirmed that the CVA20 isolate was most similar to JX275140 and KX162716 in the P2 and P3 non-structural region of the genome, respectively. Our findings therefore, show that the enigmatic non-structural region of both recombinant cVDPV2s (JX275140 and KX162716) were from indigenous NPEV-Cs. Furthermore, giving the time lapse (2008 to 2015) between isolation of both cVDPV2 isolates, our data suggests that the same indigenous population of NPEV-Cs that recombined with OPV2 to make the different lineages of cVDPV2 during the outbreak that lasted about a decade (Burns et al., 2013) might still be the ones circulating in the region.

## Conclusion

To the best of our knowledge, this is the first draft genome of CVA20 from Africa. Further, we confirm that non-reproducible CPE in L20B cell line can also be caused by enteroviruses. We had previously provided evidence (Adeniji and Faleye, 2015, Faleye and Adeniji, 2015) that the enigmatic non-structural genomic sequence in recombinant cVDPV2 in Nigeria (Burns et al., 2013) were from the indigenous circulating NPEV-Cs. Here, we confirm our previous findings unambiguously (Figures 1b and 1c). Furthermore, we show that the indigenous populations of NPEV-Cs that recombined with OPV2 to make the different lineages of recombinant cVDPV2s during the outbreak that began in 2005 were still doing the same in 2015.

Coupling the very low population immunity to the polioviruses in Northern Nigeria(Adeniji et al., 2014)with higher preponderance and circulation of NPEV-Cs in the region (Donbraye et al., 2018), we are not surprised that, as with the last cVDPV2 outbreak in Nigeria that lasted about a decade (Burns et al., 2013), the hub of the ongoing cVDPV2 outbreaks in the country is in Northern Nigeria. Hence, as long as OPV2 is in use for vaccination in the country and herd immunity to the virus is not accomplished in Northern Nigeria, recombinant cVDPV2 might remain a challenge for poliovirus eradication in Nigeria.

## CONFLICT OF INTERESTS

The authors declare that no conflict of interests exist. In addition, no information that can be used to associate the isolate analysed in this study to any individual is included in this manuscript.

### ACKNOWLEDGEMENTS

We thank the WHO National Polio Laboratory in Ibadan, Nigeria for providing the anonymouscell culture supernatant analysed in this study. This study was funded by a TETFund grant to JAA.

## DATA AVAILABILITY STATEMENT

The Sequence data that support the findings of this study are openly available in GenBank (https://www.ncbi.nlm.nih.gov/genbank/)with accession numbersMH785183 and MH785184.

